# Comparative distribution of the hypothalamic neurons activated during Wakefulness and Paradoxical (REM) sleep using TRAP2-red mice: contribution of Orexin, MCH, Lhx6 and a new marker Meis2

**DOI:** 10.1101/2025.09.17.676258

**Authors:** Amarine Chancel, Patrice Fort, Renato Maciel, Blandine Duval, Justin Malcey, Simone Bellini, Markus H. Schmidt, Pierre-Hervé Luppi

## Abstract

**Study objectives:** Paradoxical sleep (PS) is a state involving numerous hypothalamic neuronal subpopulations, many remaining neurochemically uncharacterized. Our goal was to compare hypothalamic neurons active during Wakefulness or PS rebound (PSR) and explore their potential overlap, with a focus on melanin-concentrating-hormone (MCH), Orexin (Orx), Lhx6 and a new contingent of Meis2- expressing neurons.

**Method:** In the same TRAP2-red mouse, neurons activated during Wakefulness (4h) and PSR (2h) express TdTomato and c-Fos, respectively. Double-labelling and triple immunofluorescence with neurochemical markers were performed to characterize and quantify cell populations in hypothalamic structures.

**Results:** Twelve hypothalamic structures showed distinct activation patterns. The anterior hypothalamic area (AHA), zona incerta (ZI) and tuberal nucleus contained more activated neurons during PSR than Wakefulness, whereas the paraventricular hypothalamic (PVN) and supraoptic (SO) nuclei were predominantly activated during Wakefulness. MCH and Lhx6 neurons were mainly recruited during PSR, whereas Orx neurons were activated during both. A ventral subpopulation of MCH neurons showed higher activation during PSR than the dorsal subpopulation. Additionally, ∼30% of the c-Fos+ neurons in ZI and AHA express Meis2. A similar proportion of TdTomato+ neurons positive for Meis2 were encountered in PVN and SO. Overall, ∼20% of all hypothalamic neurons activated during PSR are now neurochemically identified.

**Conclusion:** Our study identifies new neuronal populations activated during PSR in AHA and tuberal nucleus. We further get evidence that Meis2 delineates novel neuronal populations activated during PSR. In summary, our results using TRAP2-red mice characterize new cell populations activated during Wakefulness or PSR, opening experimental paths for determining their function regarding vigilance states.

**Statement of significance:** Wakefulness and paradoxical sleep are very similar at the electroencephalographic level. It remains relevant to determine the potential overlap of the neurons active during each vigilance state. We here took advantage of the powerful transgenic TRAP2-red mice to directly compare in the same animal the brain cell activation during both states, with a focus on the hypothalamus. A deeper knowledge of each individual subpopulation of hypothalamic neurons within complex brain circuits underlying the sleep-waking cycle will help the understanding and validation of treatments of sleep disorders, at least those directly linked to demonstrated hypothalamic dysfunction as Narcolepsy (Orx neurons), Amyotrophic Lateral Sclerosis (MCH and Orx signaling) or neurodegenerative diseases (Parkinson’s and Alzheimer diseases).

## Introduction

The hypothalamus is known to regulate a broad range of homeostatic functions as energy metabolism and food intake, fluid and electrolyte balance, thermoregulation, blood pressure, reproduction, circadian rhythms and vigilance states. In this regard, it contains two well-characterized wake-inducing neuronal subpopulations, the orexin/hypocretin (Orx/Hcrt) neurons in the lateral hypothalamic area (LHA) and histaminergic neurons in tuberomammillary nucleus[1–3]. Both are active during wakefulness (W) and send widespread axonal projections to the cortex, basal forebrain, limbic structures and brainstem nuclei involved in arousal [4–8]. More recently, two other populations of W-promoting neurons were discovered in LHA, i.e., glutamate CaMKIIα- expressing and GABAergic neurons which chemogenetic and optogenetic excitation induce W [9–11]. Using c-Fos or calcium imaging in mice, the paraventricular hypothalamic nucleus (PVN) was also shown to contain glutamatergic neurons active during W and inactive during sleep. Their optogenetic activation induced W while their chemogenetic inhibition or targeted-cell ablation reduced W with increase of Slow Wave Sleep (SWS) [12]. Recently, hypothalamic calretinin- glutamatergic neurons active during both W and Paradoxical Sleep (PS) were also identified in the parasubthalamic nucleus (PSTN). Similar to PVN neurons, their optogenetic and chemogenetic activation increased W and exploratory behaviours while their inhibition decreased W [13].

We previously reported in rats that, after PS hypersomnia, a very large number of neurons positive for c-Fos were localized in the posterior and tuberal hypothalamus, zona incerta (ZI), perifornical area, LHA as well as in dorsomedial (DMH) and ventromedial hypothalamic areas (VMH) [14,15]. These results highly suggest that the hypothalamus is a crucial player in PS control. Except for VMH, most of these c-Fos+ neurons express GAD67 mRNA indicating they might be GABAergic and likely inhibitory in nature [15]. We further discovered that one third of c-Fos+ GAD67- expressing neurons in LHA and ZI co-express melanin concentrating hormone (MCH) [14, 15]. Previous studies have shown that they express the enzymes responsible for GABA synthesis [16–19] while their terminals express vGAT, the vesicular GABA transporter necessary for packaging GABA into synaptic vesicles [20]. However, subsequent research contradicted these findings with several studies demonstrating that MCH neurons are not functionally GABAergic [21–25]. This was confirmed by using RNAscope and immunofluorescent staining showing that 97% of MCH neurons express vGLUT2, the vesicular glutamate transporter but not vGAT, supporting the view that MCH neurons are rather glutamatergic [26].

MCH neurons indeed fire quite specifically during PS as demonstrated by calcium imaging in mice and unit recordings in head-restrained rats [27–30]. Intracerebroventricular administration of MCH induced dose-dependent increases in PS and, to a minor extent SWS quantities in rats [14]. PS- promoting effects were partly reproduced in mice in response to optogenetic and chemogenetic activation of MCH neurons [19, 31, 32]. Interestingly, increases in PS quantities induced by thermoneutral ambient temperature warming was abolished when optogenetically inactivating MCH neurons [33]. Taken together, these data strongly support the contribution of MCH-expressing neurons in modulating PS expression.

In addition, it has been confirmed that GAD67-positive, MCH-negative, neurons in LHA and ZI discharge maximally during PS in line with our c-Fos results in rats [15, 34]. Since their increase in firing rate anticipates PS onset, such neurons could play a role in the induction of the state. We recently further showed that a subpopulation of GABAergic neurons that co-express Lhx6 in ventral ZI [35] were c-Fos+ during PS hypersomnia, forming another contingent of PS-on neurons [36]. In line with our functional data, chemogenetic activation of Lhx6+/vGAT+ neurons increases PS, whereas their inhibition suppresses it [35]. Further, calcium imaging revealed that the activity of ZI Lhx6+ neurons is maximal during PS, in line with the significant PS increase following their unilateral optogenetic stimulation [37].

We recently used TRAP2-red mice to determine the activation of MCH, Lhx6 and Orx neurons during a W period (with permanent TdTomato neuronal expression in red) followed one week apart with a PS rebound (with c-Fos neuronal expression in green) [36]. We showed for the first time that in addition to the MCH neurons, Lhx6 neurons are specifically activated during PSR while Orx ones are specifically activated during W. In that context, the aim of the present study was to extend these previous findings by determining in the TRAP2-red mice whether additional hypothalamic structures other than only the LHA and ZI may also contain neurons activated during W and/or PSR. We also re-examined the contribution of MCH-, Orx- or Lhx6-expressing neurons. In addition, a recent single-cell transcriptomic analysis revealed that molecularly distinct populations of inhibitory and excitatory neurons of LHA express the transcription factor Meis2 (Myeloid Ecotropic viral Integration Site 2, from the TALE “three amino acid loop extension” family of homeodomain-containing proteins) [24, 38]. We, therefore, studied whether Meis2 could be a new specific marker of hypothalamic neurons activated during W and/or PSR.

We first show that the anterior hypothalamic area (AHA) and Tuberal nucleus contain many neurons specifically activated during PSR. Moreover, we reveal for the first time that many Meis2- expressing neurons located in ZI and AHA are specifically activated during PSR and might play a role in PS. Overall our results indicate that around 40% of neurons activated during PSR in LHA and ZI expressed MCH, Lhx6, Orx or Meis2.

## Materials and Methods

### Animals

TRAP2-red (+/-, for targeted recombination in active populations, 2^nd^ generation) mice were generated by crossing Fos2A-iCreER/+ (TRAP2) mice (a gift by Dr. Liqun Luo, Stanford University, USA) with R26AI14/+ (AI14) mice [36, 39, 40]. TRAP2 mouse has two transgenes: one that expresses CreERT2 (i.e., tamoxifen-inducible Cre recombinase) from an activity-dependent c-Fos promoter and the other that allows expression of an effector gene (i.e., TdTomato, TdT) by means of CreERT2 recombination. With tamoxifen administration, CreERT2 recombination (i.e., binding of CreERT2 to floxed alleles) occurs, inactivating the stop codon allowing a permanent expression of TdT in trapped (activated) neurons. A sample of 8 TRAP2-red mice were used for this study. All experimental and surgical procedures were conducted in strict accordance with the European Communities Council Directive (86/609/EEC) and recommendations in the guide for the Care and Use of Laboratory Animals of the National Institutes of Health (NIH publication n° 85–23). Protocols were approved by the local CELYNE Ethics Research Committee (CEEA n° 042) of the University Claude Bernard - Lyon I and validated by the Ministère de l’Enseignement Supérieur et de la Recherche (APAFIS#21351). All efforts were made to minimize the number of animals and their suffering in agreement with the “3R” rules.

### Surgery for polysomnography

Young adult (8-12 weeks old) male TRAP2-red mice were anesthetized with Ketamine and Xylazine (100/10 mg/kg. i.p.) and placed in a stereotactic frame on a heating pad underneath at 37°C. Two home-made stainless electrodes were screwed to the skull above parietal (AP: -2.0 mm, ML: 1.5 mm from bregma) and frontal cortex (AP: +2.0 mm, ML: 1.0 mm from bregma), whereas the reference electrode for unipolar EEG recording was fixed over the occipital cortex (AP: -5 mm, ML: 0,0 mm from bregma). Two wire electrodes were inserted into the neck muscles for bipolar EMG recording. Leads were connected to a miniature plug (Plastics One, Germany) cemented to the skull. Animals recovered from surgery for at least 5 days in individual cages at constant temperature (23 ± 1 °C), humidity (30–40%) and circadian cycle (12-h light–dark cycle, lights on at 7am). Food and water were available ad libitum. Four additional TRAP2-red mice underwent the same surgery and were used for discriminating MCH subpopulations activated during PS (c-Fos+).

### Polysomnography and analysis of vigilance states

After 5 additional days of acclimatation with daily handling, prepared mice were placed in an individual Plexiglas barrel and connected with a cable to a slip-ring commutator to allow free movements during recording until sacrifice. Baseline recordings were performed the day preceding the protocol start. Unipolar EEG and bipolar EMG signals were amplified by 1:5000 V/V and 1:2000 V/V respectively (MCP-PLUS, Alpha-Omega Engineering, Israel), digitized at 1024 Hz and acquired using SlipAnalysis software (Viewpoint, France). The analysis was then performed as previously described [41, 42]. Briefly, vigilance states were scored using 5-sec window frame and classified as Waking (W), Slow Waves Sleep (SWS) or Paradoxical sleep (PS), based on the visual inspection of amplified/digitized EEG/EMG signals using SlipAnalysis. Waking was characterized by activated low-amplitude EEG accompanied by a sustained EMG activity with phasic bursts reflecting motor activity. SWS was depicted by high voltage EEG slow waves, a reduced EMG tone along with the disappearance of phasic activity. A decrease in the EEG amplitude with the distinctive theta rhythm associated with a flat EMG (muscle atonia) indicated the PS onset.

### Preparation of 4-OHT for TRAPing

4-hydroxytamoxifen (4-OHT, Cat# H6278, Sigma Aldrich, France) was prepared as previously described [36, 43]. Briefly, 4-OHT was dissolved at 20 mg/mL in absolute ethanol using an ultrasonic water bath at 37°C for 10 min, then aliquoted and stored at –20°C as a stock solution. Before using, corn oil (Sigma Aldrich) was added to the thawed stock solution to replace ethanol by evaporation at 37°C to get a 10 mg/mL concentration with a final working dose of 50 mg/kg.

### Experimental design

At 11 a.m. (ZT4), TRAP2-red mice were kept awake during 4h by placing toys and objects in their cage or gently touching them soon as they fall asleep according to EEG/EMG signals. After 2h of constant W, 4-OHT was intraperitoneally administrated, and mice were kept awake for 2 additional hours. By this way, wake-activated neurons were trapped with permanent expression of TdTomato (TdT, red fluorescence) reporter gene. One week later, the same mice were submitted to an automatic selective PS deprivation (PSD) as validated previously [41]. Briefly, each mouse was placed in an individual barrel with a movable floor connected to a small piston controlled by an electromagnet. Automatic detection of PS episodes was achieved using an algorithm previously fed with the own mouse sleep-wake features based on its baseline recording. As soon as PS is detected on-line in real time, a TTL (transistor-transistor logic) pulse was send to the electromagnet, causing brief floor movements (up/down) that awoke the mouse within 5 sec. The PSD started at 11 a.m. for 48h. Mice were then allowed to recover their sleep debt for 2h (PSR) before sacrifice. The PSR started with the first occurrence of a PS episode (Figure 1A).

**Figure 1.**
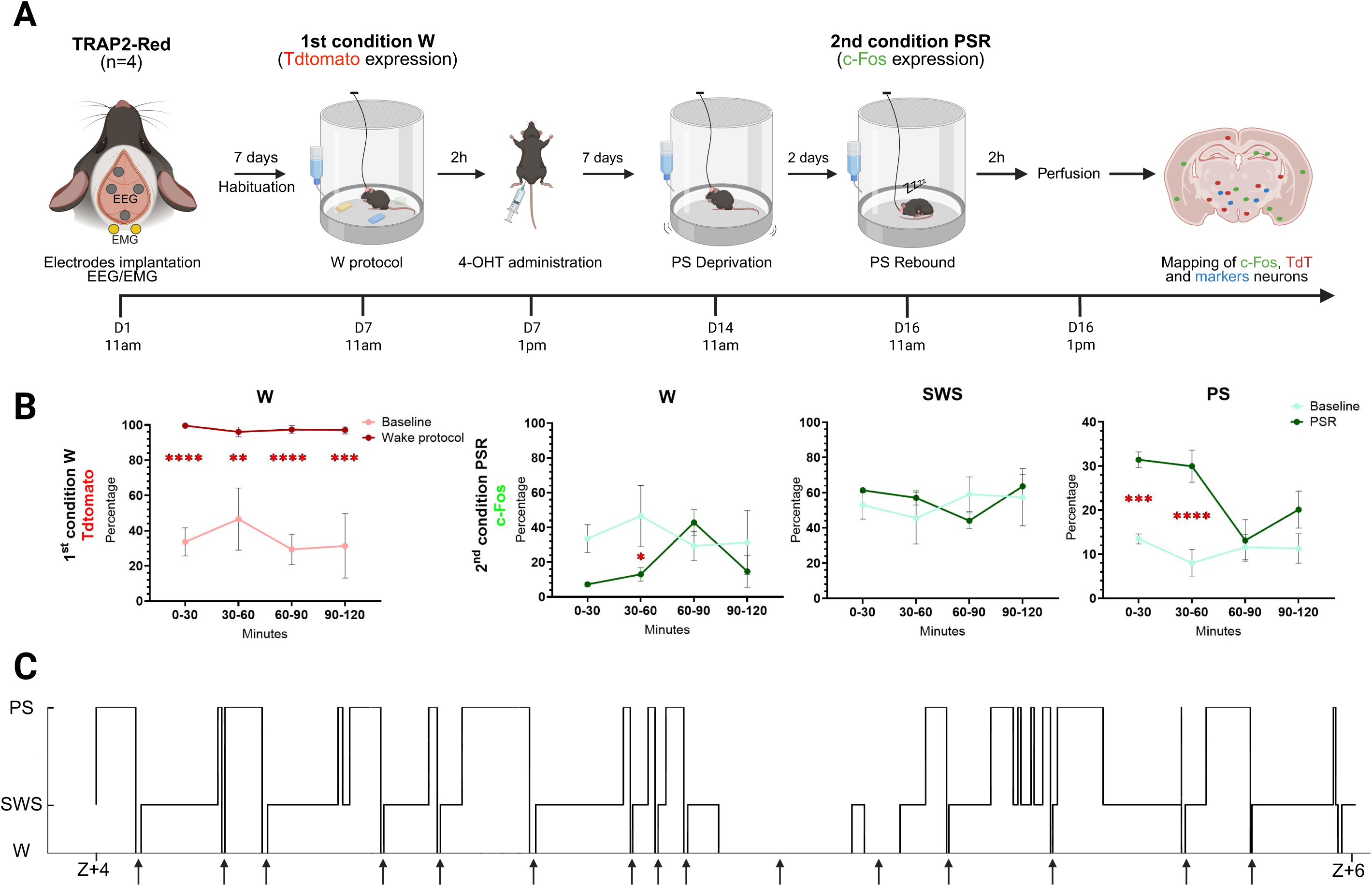
Experimental design. **(A)** Time course of experiments in TRAP2-red mice. After 2h of W, 50mg/kg of 4-OHT was injected I.P. to trap neurons activated during the W protocol (TdT+). One week later, the same mice were exposed to 48h of PSD followed by 2h of PSR (c-Fos+) before sacrifice. Then, mapping of neurons positive for c-Fos, TdT and other markers of interest were done within the whole hypothalamus. **(B)** Comparison of mean W percentage between baseline and W protocol (left panel). On right panel, comparison of mean W, SWS and PS percentages during PSR compared to baseline (from 11am to1pm.). Shapiro Wilk and a two-way ANOVA with Tukey’s multiple comparisons post-hoc test were applied in the case of significance (****p < 0.0001; ***p < 0.001; **p < 0.01; *p < 0.05, n=4). **(C)** Hypnogram from a representative TRAP2-red mouse illustrating the expected large amounts of PS during the 2h PSR preceding sacrifice. Noticed the high number of short bouts of W (arrows). W: wakefulness; PSR: paradoxical sleep rebound; SWS: slow-wave sleep; PS: paradoxical sleep; EEG: electroencephalogram; EMG: electromyogram

### Histological procedure

At 1 p.m. (ZT6), mice were anesthetized (pentobarbital 140 mg/kg, I.P.) and transcardially perfused with heparinized Ringer’s lactate solution, followed by 4% paraformaldehyde (PFA) in phosphate-buffered saline (PBS, pH 7.4). Brains were removed, post-fixed overnight in 4% PFA at 4°C and stored in 30% sucrose PBS solution at 4°C for at least 48h. They were frozen at -40°C in methylbutane and cut into 30 µm thick coronal sections using a cryostat. Sections were collected in 8 wells containing cryoprotective solution (PBS 0.05 M, 20% glycerol, 30% ethylene glycol, pH 7.4) and stored at -20°C.

### Immunofluorescence

After initial washing in 0.1 M PBS with 0.3% Triton X-100 (PBS-T, Sigma-Aldrich), free-floating sections were then treated for multiplex immunofluorescent staining. They were rinsed in 0.3% H2O2 solution in PBS-T for 1h to quench endogenous peroxidase activity. After 3 washes, they were incubated in primary antibodies against c-Fos (monoclonal rat, 226 017, 1:50000, Synaptic System, Germany) and either MCH (polyclonal rabbit, H-070-47, 1:20000, Phoenix Pharmaceutical, USA), Lhx6 (monoclonal mouse, sc-271433, 1:1000, Santa Cruz, USA), Meis2 (polyclonal rabbit, NBP1-81669, 1:5000, Novusbio, UK) or Orexin (polyclonal goat, sc-80263, 1:10000, Santa Cruz) diluted in a PBS-T containing sodium azide (0,1%) for 2-3 days at 4°C under constant stirring.

After washing, sections were then incubated for 3h at room temperature in PBS-T containing biotinylated donkey anti-rat IgG antibody (1:1000, Vector Laboratories, USA) for c-Fos detection and either donkey anti-rabbit IgG-alexa 647 antibody (InvitroGene, 1:500) for MCH or Meis2, donkey anti-mouse IgG-alexa 647 antibody (InvitroGene, 1:500) for Lhx6 or donkey anti-goat IgG-alexa 647 antibody (Invitrogen, A21447, 1:500) for Orexin. After rinses sections were incubated with ABC-HRP solution (1:1000; Vectastain Elite kit, Vector Labs) for 90 min at room temperature. After several washes in PBS-T, they were revealed for 10 min in Alexa 488-conjugated tyramide (Tyramide SuperBoost™ Kit; InvitroGene, Life Technologies, Eugene, USA, B40953, 1:500). The reaction was terminated with several rinses with PBS-T. Immunoreacted sections were mounted on gelatine-coated glass slides, dried and finally cover-slipped with prolong Gold anti-fade reagent containing 4’,6-diamidino-2-phenylindole (DAPI) (Molecular Probes, Eugene, OR, USA) and stored at 4°C.

### Cell counting

Immunoreacted hypothalamic sections were imaged using an Axioscan Z1 scanner (Zeiss, Germany). Images were collected with a x20 objective (N.A. Plan-Apochromat 20x/0.8 M27 Air/0.8.) and a 0.45 Orca Flash camera. DAPI (405, blue), c-Fos (PSR-activated neurons, 488, green), TdTomato (W-trapped neurons, 555, red-orange), MCH, Lhx6, Meis2 or Orx (647, deep red) were acquired using appropriate filter cubes according to manufacturer recommendations (CIQLE, Lyon-Est Centre for Quantitative Imaging, Lyon I University). For scanned sections, five mosaics were acquired (5µm steps) then stacked to a single plane, using the depth of “focus wavelet” function thanks to Zen software (version 3.1, Zeiss). Images were directly imported into NeuroInfo software (Version 2023 1.5, MBF Bioscience, USA) allowing automatic detection and quantification of fluorescent objects at different wavelengths associated to hypothalamic areas identified by matching sections to a three-dimensional reference atlas (Third Allen Atlas, Brain Common Coordinates Framework CCF v3). A manual correction was required to align boundaries of each hypothalamic nucleus to each section for each mouse. (n=4). Drawings of structures and plotting of c-Fos+ (green) and TdT+ (red-orange) neurons were done in addition to one marker, i.e., MCH, Lhx6 Meis2 or Orx (deep red). Putative double- (c-Fos+/TdT+, c-Fos+/marker+, TdT+/marker+) and triple-labelled (c-Fos+/TdT+/marker+) neurons were also automatically encountered.

For each mouse, 8 hypothalamic sections separated by 240µm from Bregma -0.45 to 2.20 were analysed. Counts for each type of labelled neurons for a structure found on several successive sections were summed: LHA (found on 8 sections); ZI (7 sections); AHA and Arc (5 sections); VMH and PVN (4 sections); DMH, SO, Tub and RCh (3 sections); PSTN and PHN (2 sections). As no statistical difference was found between the two hemispheres, counting from both sides were summed.

The percentage of double-labelled neurons during PSR and W was obtained by dividing the number of c-Fos+-TdT+ by the total number of c-Fos+ or TdT+ neurons respectively, for each structure. The closer the percentage is to zero for a structure, the less the structure is reactivated in W or PS. On the contrary, the higher the ratio, the more neurons are reactivated in both conditions (Figures 2B).

**Figure 2.**
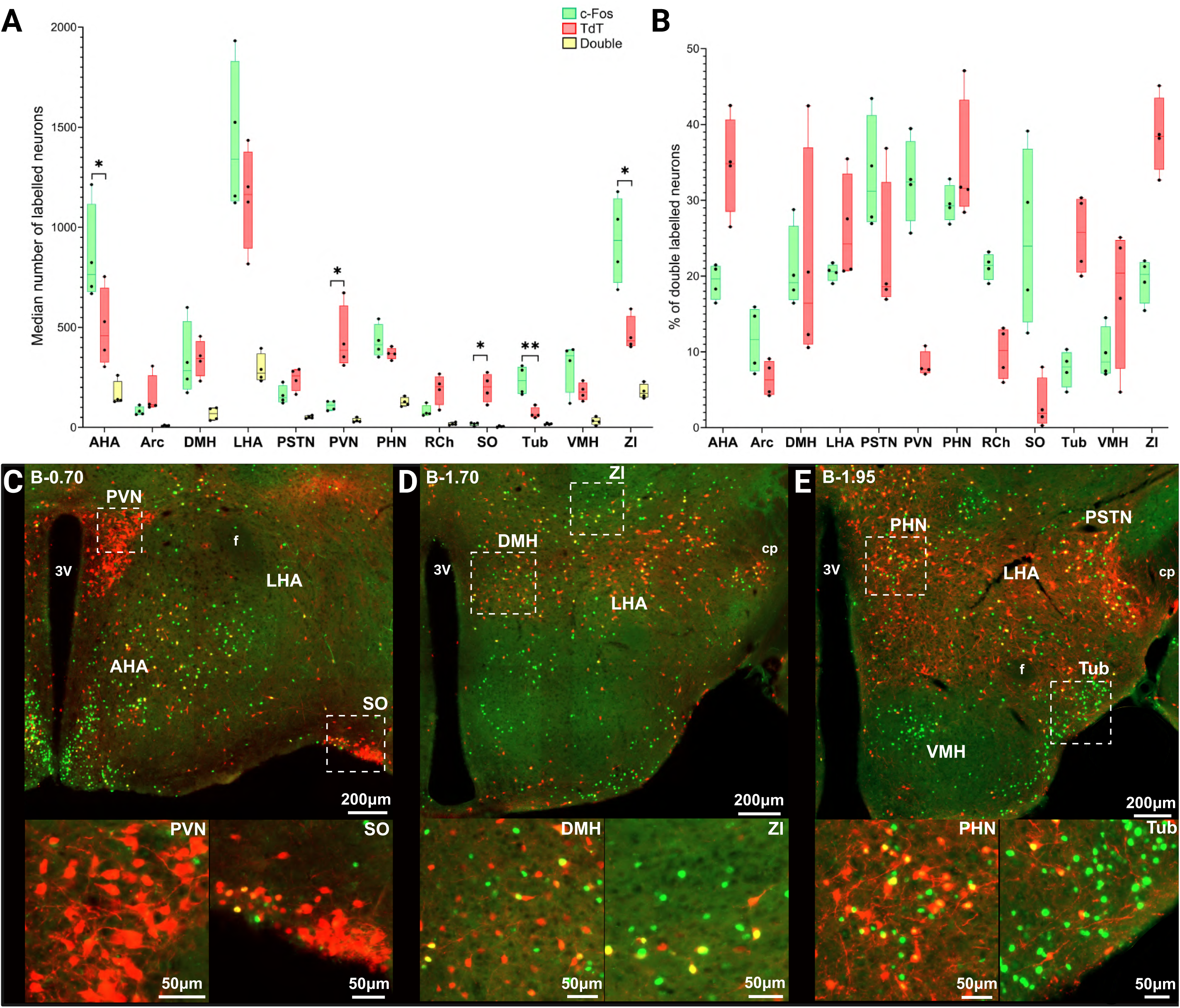
Comparative distribution and quantification of c-Fos+, TdT+ and double-labelled neurons in the 12 hypothalamic structures of interest. **(A)** Comparison of the median number of c-Fos+ and TdT+ and double-labelled neurons in 12 hypothalamic structures. AHA, ZI (*p<0.05) and Tub (**p<0.01) contain significatively more c-Fos+ than TdT+ neurons. PVN, RCh and SO (*p<0.05) contain significatively more TdT+ than c-Fos+ neurons (n=4 mice). **(B)** Percentage of double-labelled neurons over the total number of c-Fos+ neurons (green) and over the total number of Tdt+ neurons (red) in each hypothalamic structure, n=4. **(C)** Low- and high-power photomicrographs at Bregma-0.70 showing the PVN and SO which contain mostly TdT+ neurons (red) and a few c-Fos+ neurons (green). The yellow neurons are double labelled with c-Fos and TdT. Bars: 200 and 50µm, respectively. **(D)** Low- and high-power photomicrographs at Bregma -1.70 showing DMH which contains a substantial number both of c-Fos+ (green) and TdT+ (red) neurons and ZI with more c-Fos+ (green) than TdT+ neurons (red). Both nuclei contained a small proportion of double-labelled neurons. Bars: 200 and 50µm, respectively. **(E)** Low- and high-power photomicrographs at Bregma-1.95 showing the PHN which contains many c- Fos+ (green) neurons, TdT+ (red) and double-labelled (yellow) neurons and the Tub containing many more c-Fos+ than TdT+ neurons. Bars: 200 and 50µm, respectively. AHA: anterior hypothalamus nucleus; Arc: arcuate nucleus; DMH: dorsomedial hypothalamus nucleus; LHA: lateral hypothalamus area; PHN: posterior hypothalamic nucleus; PSTN: parasubthalamic nucleus; PVN: paraventricular hypothalamic nucleus; RCh: retrochiasmatic area; SO: supraoptic nucleus; Tub: tuberal nucleus; VMH: ventromedial hypothalamus; ZI: zona incerta.

### Statistics

For polysomnographic and neuroanatomical data, statistics were conducted using GraphPad Prism 10.4.2. They were expressed as median ± SEM. Normality was tested with a Shapiro-Wilk Test. Statistical significance was defined at a threshold of 5% (p<0.05). p-Values are indicated in the text and captions of Figures. For wake-sleep data, a two-way analysis of variance (ANOVA) with Tukey’s multiple comparisons post-hoc test was applied in the case of significance (Figure 1B). Paired t-test were conducted to compare median number of labelled neurons, percentage of double- and triple-stained neurons in each hypothalamic structure (Figures 2A; 3A, B, E; 4C-D).

**Figure 3.**
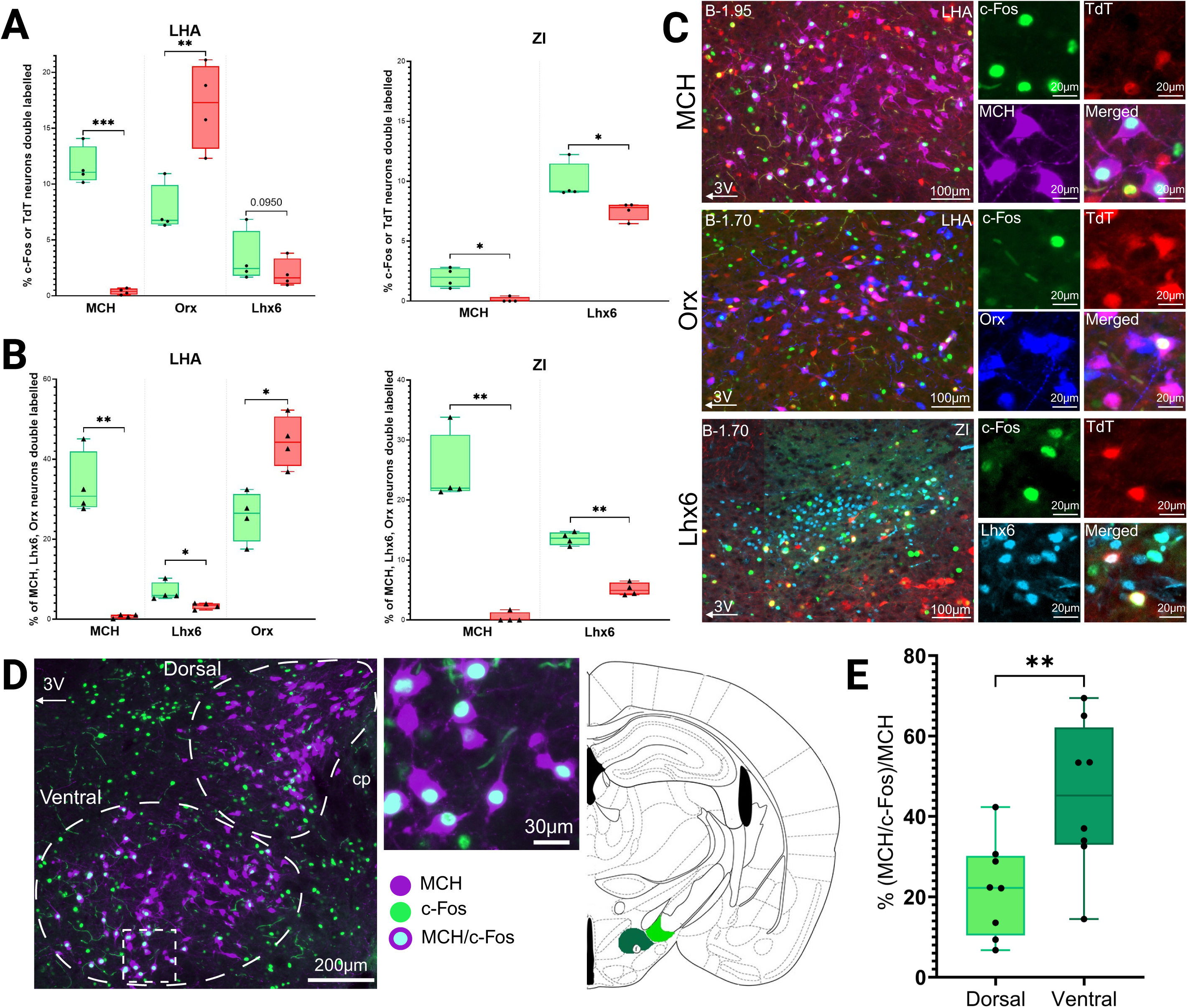
Distribution of c-Fos+ and TdT+ double-labelled neurons with either MCH, Orx or Lhx6 in LHA and ZI. **(A)** Bar plots showing the percentage of c-Fos+ (green) and TdT+ (red) neurons that are also positive for either MCH, Orx or Lhx6 in LHA and ZI. Significant differences between c-Fos+ and TdT+ neurons were found in LHA for MCH+ (***p<0.0007) and Orx+ (**p<0.0074), but not for Lhx6+ neurons (p = 0.095). In ZI, significant differences are found for c-Fos+ and TdT+ also positive for MCH (*p<0.0161) or Lhx6 (*p<0.0430). Bar plots showing the percentage of MCH, Orx, or Lhx6 neurons that are c-Fos+ (green) or TdT+ (red) in the LHA and ZI. Significant differences between c-Fos+ and TdT+ neurons were found in the LHA for MCH (**p < 0.0034), Lhx6 (*p <0.0439) and Orx staining (*p<0. 0.0355), as well as in ZI for MCH (**p<0.0047) and Lhx6 (**p<0.0032) labelling. **(B)** Low- and high-power photomicrographs showing MCH+, Orx+ and Lhx6+ neurons also positive for c-Fos or TdT or both in LHA and ZI. Bars = 100 and 20µm, respectively. **(C)** Left: Representative photomicrographs (bars = 200 µm and 30µm) showing MCH neurons (purple) with (white nucleus) or without c-Fos staining in LHA. Right: drawing from a mouse brain atlas (Bregma –1.95 mm) showing the localization of the two MCH subpopulations — one dorso-medial to the cerebral peduncle (cp, MCHd, light green) and the other, with a ventral position close to the third ventricle (3V, MCHv, dark green) — More c-Fos+ neurons are visible in the ventromedial group. **(D)** Percentage of MCH+ neurons expressing c-Fos in the dorsal (light green) and ventral (dark green) subpopulations during PSR. The ventral one shows a higher level of activation (**p < 0.0016). *n = 8 animals*. All statistics: Shapiro-Wilk normality test and paired t-test; data presented as median ± SEM LHA: lateral hypothalamus area; Lhx6: Lim homeobox 6; MCH: melanin-concentrating hormone; Meis2: Myeloid Ecotropic viral Integration Site 2 homeobox; Orx: orexin/hypocretin; TdT: TdTomato; ZI: zona incerta.

**Figure 4.**
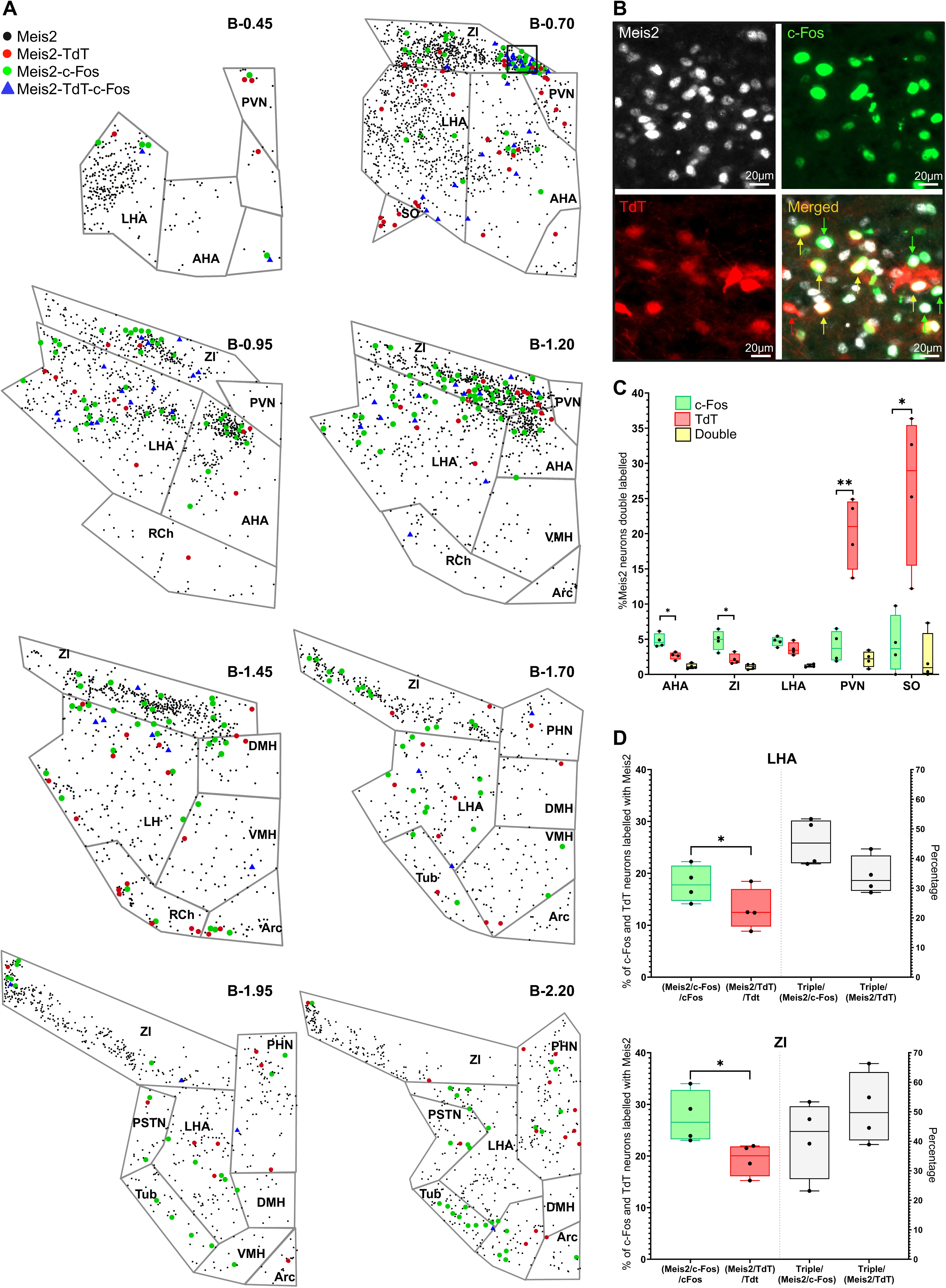
Distribution and quantification of Meis2+, Meis2+/c-Fos+, Meis2+/TdT+ and triple-labelled neurons. **(A)** Cartography of hypothalamic Meis2+ (black dot), Meis2+/c-Fos+ (green dot), Meis2+/TdT+ (red dot) and Meis2+/c-Fos+/TdT+ (blue triangle) neurons from B-0.45 to B-2.20 in a representative TRAP2- red mouse. **(B)** Photomicrographs illustrating Meis2+ (white), c-Fos+ (green), TdT+ (red) and double- (green arrow) and triple-labelled (yellow arrow) neurons within ZI at B-0.70. Scale bars: 20µm. **(C)** Histograms showing the percentage of Meis2+ neurons double- or triple-labelled with c-Fos and TdT. For AHA, Meis2+/c-Fos+ neurons are significantly more numerous that Meis2+/TdT+ ones (*, p< 0.0177). Similar results for ZI (*, p< 0.0489). For PVN, Meis2+/TdT+ neurons are significantly more numerous that Meis2/c-Fos+ ones (**, p< 0.0100). Similar results for RCh and SO (*, p< 0.0201). For statistics Shapiro Wilk and a paired T-test were applied, median ± SEM, n=4. **(D)** Percentage of c-Fos+ and TdT+ neurons that are Meis2+ and percentage of triple-labelled neurons in LHA and ZI. Significant differences are observed between the percentage of Meis2+/c-Fos+ neurons and Meis2+/TdT+ neurons in LHA (*p<0.0347) and ZI (p*<0.0135). Gray plots on the right display the percentage of triple-labelled neurons relative to the total number of c-Fos+ or TdT+ neurons. AHA: anterior hypothalamus nucleus; Arc: arcuate nucleus; DMH: dorsomedial hypothalamus nucleus; LHA: lateral hypothalamus area; Meis2: Myeloid Ecotropic viral Integration Site 2 homeobox; PHN: posterior hypothalamic nucleus; PSTN: parasubthalamic nucleus; PVN: paraventricular hypothalamic nucleus; RCh: retrochiasmatic area; SO: supraoptic nucleus; TdT: TdTomato; Tub: tuberal nucleus; VMH: ventromedial hypothalamus; ZI: zona incerta.

## Results

### Analysis of the vigilance states

Wakefulness (W) was induced at 11 a.m. using sensory stimulation for 4h. After 2h, 4-OHT was I.P. administrated and mice were maintained awake for 2 additional hours (Figure 1A). They displayed almost 100% W during the first 2h protocol with highly significant differences in hourly W percentages compared to baseline (Figure 1B). One week later, a PS rebound (PSR) was induced in response to 48h of selective PS deprivation. During the 2h preceding sacrifice, PS quantities significantly increased compared to baseline during the 1^st^ (***p<0.0007) and 2^nd^ half-hour (****p<0.00001) (Figure 1B, C). Such increases were at the expense of W, while SWS quantities remained unchanged (Figure 1B). Short bouts of W (number: 22 ± 1.8; mean duration: 16.5±5.4 s) occurred during PSR.

### Mapping and quantification of c-Fos+, TdT+ and double-labelled neurons in the hypothalamus

Cell counting was done for 12 hypothalamic structures (mean cell numbers and densities) to evaluate their activation level during PSR and W (Figure 2A and Figure Supplementary 1). These areas were the anterior hypothalamic area (AHA), arcuate hypothalamic nucleus (Arc), dorsomedial nucleus of the hypothalamus (DMH), lateral hypothalamic area (LHA), parasubthalamic nucleus (PSTN), paraventricular hypothalamic nucleus (PVN), posterior hypothalamic nucleus (PHN), retrochiasmatic area (Rch), supraoptic nucleus (SO), tuberal nucleus (Tub), ventromedial hypothalamic nucleus (VMH) and zona incerta (ZI).

### Distribution of c-Fos+ and TdT+ hypothalamic neurons

As illustrated in Figure 2A, the LHA contained the largest number of c-Fos+ and TdT+ neurons, followed by ZI and AHA. Three among 12 hypothalamic structures contained significantly more c-Fos+ than TdT+ neurons, i.e., AHA (*p<0.0167), ZI (*p<0.0246) and Tub (**p<0.0095). Two structures depicted significantly higher numbers of TdT+ than c-Fos+ neurons, i.e., SO (*p<0.0142) and PVN (*p<0.0236). No significant difference was found for the remaining 7 structures (Fig. 2A).

### Percentage of c-Fos+/TdT+ double-labelled neurons

When considering the whole hypothalamus, less than 20% of c-Fos+ neurons were also labelled for TdT, indicating that 80% of c-Fos+ neurons are selectively recruited during PSR (Fig. 2A and B). Conversely, less than 20% of TdT+ neurons were also c-Fos+, indicating that 80% of TdT+ neurons were activated specifically during W (Figure 2B). Four types of hypothalamic structures were discriminated based on their activation pattern.

The first category was composed of structures containing significantly more neurons activated during PSR than W and with a low percentage of double-labelled neurons, i.e., ZI, AHA and Tub (Figure 2). The ZI contained the highest number of c-Fos+ neurons, which was significantly higher than for TdT+ (*p<0.0246 median number). Further, 80% of c-Fos+ neurons and 62% of TdT+ neurons were not double-labelled. The AHA was the second structure with significantly more c-Fos+ than TdT+ neurons (*p<0.0167). As for ZI, 80 % of c-Fos+ neurons were activated only during PSR while 65% of TdT+ neurons were activated only during W. Finally, the Tub also contained significantly more c-Fos+ than TdT+ neurons (**p< 0.0095 median number cells and *p< 0.0188 density cells). Interestingly, 92% of c-Fos+ neurons and 74% of TdT+ cells were not double-labelled indicating that this nucleus is very specifically activated during PSR.

The second category includes structures containing mostly neurons activated during W, i.e., PVN, SO and RCh. Regarding PVN, it essentially contained many packed TdT+ neurons and only a few c-Fos+ neurons (*p<0.0236 median number; Figure 2). Of these TdT+ neurons, 92% were not double-labelled, while on the contrary, 32% of c-Fos+ neurons were also activated during W. The SO and RCh displayed the same profile (*p<0.0142 and p=0.0536 median number of cells, respectively). For SO, 98% of TdT+ neurons and 76% of c-Fos+ neurons were not double-labelled. Regarding RCh, 90% of TdT and 79% of c-Fos+ neurons were not double-labelled.

The third category of structures, i.e. Arc, VMH and DMH, showed a similar number c-Fos+ and TdT+ neurons, and low percentage of double-labelled neurons (between 10 and 20%). No significant difference in the number of c-Fos+ and TdT+ neurons was observed in Arc (p= 0.2309 median number). Interestingly, 88% of c-Fos+ neurons and 94% of TdT+ neurons were not double-labelled indicating that Arc contains two distinct cell populations specifically activated during W or PSR. The same is true for VMH which contained similar number of c-Fos+ and TdT+ neurons (p=0.1588 median number) mostly not double-labelled (91% for c-Fos+ and 80% for TdT+ neurons). The same results were observed for DMH, with no significant difference and a quite large number of c-Fos+ and TdT+ neurons (p=0.9475 median number) mostly not double-labelled (81% for c-Fos+ and 84% for TdT+ neurons) (Figure 2).

The fourth category of structures contained a large (LHA) and a substantial (PSTN, PHN) number of TdT+ and c-Fos+ neurons with a percentage of double-labelled neurons superior to 20% (Figure 2A, B). The LHA showed the largest number of c-Fos+ and TdT+ neurons (p=0.2191). Of interest, 79% of c-Fos+ and 76% of TdT+ neurons were not double-labelled indicating that LHA contains two contingents of neurons specifically activated during W or PSR. The PSTN and PHN contained a significant number of TdT+ and c-Fos+ neurons in both conditions (p=0.2121 and p=0.3095, respectively). In these areas, a high proportion of the c-Fos+ neurons (70 and 71%, respectively) and TdT+ cells (81 and 68%, respectively) neurons were not double-labelled (Figure 2).

### Activation levels of MCH, Lhx6 and Orx neurons during W or PSR

In LHA, 11 ± 0.9% of c-Fos+ and 0.4 ± 0.1% of TdT+ neurons co-expressed MCH. Additionally, 6.8 ±1.1% of c-Fos+ and 17.3 ±1.9 of TdT+ neurons were positive for Orx (**, p< 0.0074). Finally, 2.4 % of c-Fos+ and 1.6% of TdT+ neurons were positive for Lhx6 (p=0.0950). Since it has previously been shown that the three populations are different, it can be concluded that about 20% of neurons in LHA activated during PSR and 18% of those recruited during W are neurochemically identified (Figure 3A, C).

In ZI, 2 ± 0.4% of c-Fos+ and 0 ± 0.1% of TdT+ neurons were MCH positive (*, p< 0.0161). In sharp contrast, 9.2 ± 0.8% of c-Fos+ and 7.8 ± 0.4% of TdT+ neurons were Lhx6+ (*p<0.0430). These results suggest that more than 90% of neurons activated during PSR or W in the ZI are still uncharacterized (Figure 3A, C).

Importantly, 30.8 ± 4.0% and 22 ± 3.0% of MCH+ neurons in LHA and ZI were positive for c-Fos while nearly none were TdT+ (**p < 0.0034 and **p<0.0047 respectively) or triple-labelled (0.3% in LHA; 0.2 % in ZI). Concerning Lhx6 neurons, 6 ±1.1% of them in LHA and 13.6 ± 0.5 in ZI were c-Fos+, while only 3.5 ± 0.4% in LHA and 4.9 ± 0.5% in ZI were TdT+ (*p < 0.0439 and **p < 0.0032, respectively; Figure 3B, C). A few of them were occasionally triple labelled (1.6% in LHA and 3% in ZI). Finally, a large percentage of Orx+ neurons were activated during W (44.2 ±3.2%). Surprisingly, 26.5 ±3.1% of Orx+ neurons were also c-Fos + in PSR condition (*p<0.0355) and 11.4% were activated both during W and PSR.

Finally, we found that two different subpopulations of MCH+ neurons can be functionally distinguished in the caudal LHA (from Bregma -1.70) (Fig. 3D). At that level, MCH neurons could be separated into a ventromedial subpopulation containing significantly fewer MCH+ neurons than in the dorsolateral subpopulation (median MCHd = 313.5 vs MCHv = 188). Interestingly, a significantly larger proportion of MCH+ neurons were c-Fos+ in the ventromedial than in the dorsolateral subpopulation (45.2 ±6.6% vs 22.2 ± 4.2%; **p<0.0016) (Figure 3D, E).

### Meis2: a new hypothalamic marker of interest

Based on Mickelsen et al. [24], we further tested whether Meis2 could be a marker for uncharacterized hypothalamic neurons activated during PSR and/or W. A very high number of Meis2-immunoreactive neurons were localized in ZI (median number = 4412.5; Figure 4B), LHA (4087.5) and to a minor extent in AHA (1171.0) (Figure 4A). Interestingly, a much higher number of Meis2+ neurons were localized in the rostral than caudal ZI. A much smaller number of Meis2+ neurons was distributed in other hypothalamic nuclei (Figure 4A, Table 1).

**Table 1.** Quantification of c-Fos+, TdT+, Meis2+, Meis2+/c-Fos+, Meis2+/TdT+ and Meis2+/c- Fos+/TdT+ neurons in each hypothalamic structure of interest. Eight coronal sections were counted for each animal (n=4). Data are presented as median ± SEM. AHA: anterior hypothalamus nucleus; Arc: arcuate nucleus; DMH: dorsomedial hypothalamus nucleus; LHA: lateral hypothalamus area; Meis2: Myeloid Ecotropic viral Integration Site 2 homeobox; PHN: posterior hypothalamic nucleus; PSTN: parasubthalamic nucleus; PVN: paraventricular hypothalamic nucleus; RCh: retrochiasmatic area; SO: supraoptic nucleus; TdT: TdTomato; Tub: tuberal nucleus; VMH: ventromedial hypothalamus; ZI: zona incerta.

| Mean±SEM | Section nbr | c-Fos | TdT | Meis2 | Meis2/c-Fos | Meis2/TdT | Meis2/c-Fos/TdT |
| --- | --- | --- | --- | --- | --- | --- | --- |
| AHA | 5 | 661,3±97,2 | 471±80,9 | 1171±108,9 | 56,5±7,9 | 31±4,3 | 13,8±2,8 |
| Arc | 5 | 50,8±14,3 | 170,8±43,7 | 116,3±37,6 | 4,8±3,1 | 9±2,7 | 0,0±0,0 |
| DMH | 3 | 250,3±53 | 378±60,8 | 141±26,7 | 5,5±1,7 | 5±0,7 | 0,3±0,3 |
| LHA | 8 | 1070,8±125,7 | 1136,3±107,7 | 4087,5±529 | 195,8±38,3 | 142,8±11 | 49±6,5 |
| PSTN | 2 | 182±7,5 | 314,8±79,9 | 109,8±25,9 | 4,8±1,8 | 4,3±2,4 | 0,8±0,5 |
| PVN | 4 | 94,3±22 | 438,8±71,3 | 173,5±35 | 8±3,5 | 35,3±9,9 | 4,3±1,8 |
| PHN | 2 | 454,5±57 | 434,8±50,7 | 376±95,9 | 18,5±5,6 | 12,5±3,9 | 5±1,4 |
| RCh | 3 | 73±16,5 | 249,5±25,5 | 166,8±35,6 | 4,3±1,2 | 24,3±6,9 | 1,8±0,3 |
| SO | 3 | 16,8±5 | 195,8±31,5 | 116,3±46,9 | 3,5±1,4 | 34,5±16,6 | 1,3±0,6 |
| Tub | 3 | 198,5±39,9 | 61±10,5 | 103±26,3 | 7,3±3 | 1,8±1,0 | 0,3±0,3 |
| VMH | 4 | 211±50,8 | 171±30,6 | 180,3±49,6 | 5,8±3,1 | 3,5±0,9 | 0,8±0,3 |
| ZI | 7 | 807±134,4 | 500,8±66,2 | 4512,5±301 | 224,8±45,8 | 95,8±12,3 | 49,3±8,4 |
| Total | 8 | 4575±487,7 | 4736,3±468,4 | 11326±1164,4 | 546±103,6 | 403,3±29,9 | 127,5±15,9 |

Many Meis2+ neurons expressed c-Fos in ZI (224.8±45.8), LHA (195.8±38.3) and to a minor extent AHA (56.5±7.9), whereas other hypothalamic nuclei contained a small number of c-Fos+/Meis2+ neurons (Table 1). There were approximately two-times the number of c-Fos+/Meis2+ than TdT+/Meis2+ neurons in ZI (* p< 0.0489) and AHA (* p< 0.0177) but not in LHA (p=0.1581) (Table 1). In the latter structure, 17.8% of c-Fos+ and 12.5% of TdT+ neurons were Meis2+ (*p<0.0347). In ZI, 26.5% of c-Fos+ and 20% of TdT+ neurons were Meis2+ (p*<0.0135). However, in all structures, c-Fos+/Meis2+ and Tdt+/Meis2+ neurons constituted a small percentage of the total number of Meis2+ neurons (ZI, c-Fos+, 5 ± 0.7%; TdT+, 2 ± 0.4%; LHA, c-Fos+, 4.8 ±0.3%, TdT+, 3.4 ± 0.5%; AHA, c-Fos+, 4.5 ± 0.5; TdT, 2.7 ± 0.3%) (Figure 4A and C). We further determined whether TdT+/Meis2+ and c-Fos+/Meis2+ neurons were belonging to the same population. Approximately one fourth of c-Fos+ neurons in LHA (25.8 ± 2.3%) and ZI (24.8 ± 3.7%) were indeed triple labelled (Fig 4D). One third of TdT+ neurons in LHA (32.6 ± 3.2%) and half in ZI (49.7 ± 6.0%) were triple labelled. These results indicate that most of LHA and ZI Meis2+ neurons activated during PSR were not activated during W, in contrast to up to half of neurons activated during W that were also activated during PSR.

Finally, the PVN and SO contained a significant number of TdT+/Meis2+ neurons but only a few c-Fos+/Meis2+ cells (Table 1). Further, a significantly higher percentage of TdT+ neurons expressed Meis2 (21 ± 2.6% and 29 ± 5.3%, respectively) compared to c-Fos+ (3.7 ±1.1% and 3.7% ± 2.1%; ** p< 0.0100 and * p< 0.0458, respectively).

## Discussion

In the present study, the methodological advances brought by the transgenic TRAP2-red mice were used to map the neurons sequentially activated one week apart during W (first step, TdT expression) and then PSR (second step, c-Fos immunodetection) in 12 hypothalamic nuclei. We first re-examined and compared the levels of activation of MCH+, Orx+ and Lhx6+ neurons in LHA and ZI during W or PSR. Then, we showed that additional structures such as the Tub, AHA, PVN and SO are specifically activated during W or PSR. Finally, we showed for the first time that Meis2 is a new marker of PSR neurons not expressing MCH, Orx or Lhx6 in AHA, LH and ZI.

The hypothalamus is a highly heterogeneous area in terms of its neuronal populations and functions. Our results indicate that most structures among those analysed, excepting PVN and SO, contain neurons activated during W or PSR. We further showed that around 20% of the LHA-ZI neurons express c-Fos and TdT and are, therefore, activated during both conditions. Alam et al., [44] previously showed that more than half of the perifornical neurons within LHA exhibited peak discharge rates during W and PS. About 38% of neurons were classified as W-active with decreased firing rates both during SWS and PS. In contrast, ZI contained significantly more neurons activated during PSR than W in agreement with our previous c-Fos studies in rats [14, 15] and more recently in TRAP2-red mice [36]. GABAergic neurons were recorded in ZI by electrophysiology and calcium imaging, and more than half of these neurons increased their activity during PS [45, 46]. In our experiment, GABAergic Lhx6 neurons are labelled with c-Fos during PSR after 48h of PS deprivation but not during W. In contrast, Liu et al., (2017) and Lee et al., (2025) [35, 47] showed that during PSD, glutamatergic Reuniens neurons activate Lhx6+ GABAergic neurons in the ZI. Their activation may prepare or promote the rebound of PS that follows 72h of PSD. The authors proposed that Lhx6 neurons inhibit W systems such as Orx neurons [35]. Our results and those of others do not support their hypothesis of an activation of Lhx6 neurons during sleep deprivation and in contrast indicate that they are mostly activated during PS. Additional data is necessary to resolve these contrasting findings.

In addition, we report for the first time that AHA and Tub contain higher numbers of neurons activated during PSR than during W and that only a few cells were activated during both states. These data are in line with our previous study in rats showing a strong activation of tuberal neurons during PSR [15]. To our knowledge, the AHA has not been studied in the context of sleep. The identification of a specific marker for these neurons is now required to better characterize their activity and manipulate them with opto- or chemogenetic approaches.

The other hypothalamic structures such as PSTN, DMH, Arc, VMH and PHN contain similar numbers of c-Fos+ and TdT+ neurons. Around 25% of these labelled neurons were activated both during W and PSR meaning that approximately 75% are selective for either state. Supporting our data, glutamatergic PSTN neurons were shown to be active during W and PS and silent during SWS [13]. The opto- and chemogenetic activation of these neurons or their efferent inputs to ventral tegmental areas and lateral parabrachial nucleus, increases W and exploratory behaviours while their inhibition increases sleep. Regarding DMH, it is known to relay circadian signals from the suprachiasmatic nucleus (SCN) to sleep–wake circuits, modulating both sleep-promoting neurons in ventrolateral preoptic nucleus (VLPO) and main arousal-promoting signalling systems such as Orx-, histamine- or noradrenaline-releasing neurons [48]. GABAergic neurons in DMH promote W in response to cold exposure and facilitate recovery from anaesthesia [49, 50]. A distinct W-active subpopulation within DMH in mouse, acts as a circadian regulator, preventing excessive arousal during the night [51]. In addition, the DMH also contain two subpopulations of neurons active during PS (PS-on) projecting to the raphe pallidus or inactive during it (PS-off) projecting to the preoptic area. The activation of the DMH PS-on neurons increases PS amounts, while the activation of PS-off neurons decreases PS [52]. The arcuate nucleus is well known to be involved in metabolism with NPY/AgRP and POMC/CART neurons, and our results suggest that their activity might be increased during W or PS [53–57]. Regarding VMH, an early lesion study in rats showed that cell loss disrupts normal sleep–wake patterns along with altered feeding behaviours and metabolic dysregulation[58]. On the contrary, Hirasawa et al., [59] identified a specific subpopulation of VMH neurons depicting their highest activity during PS. Lastly, the posterior hypothalamus (PH) is a key wake-promoting centre as its lesion in cats and rats causes profound hypersomnia [60, 61]. Unit recordings in freely moving cats showed that PH contains neurons with sustained firing during both W and PS, suggesting their role in cortical desynchronization characterizing both vigilance states [62]. In line with those data, PSTN, DMH, Arc, VMH and PHN indeed contains a substantial number of c-Fos+/TdT+ double-labelled neurons in our experiments in TRAP2-red mice.

Finally, the PVN and SO predominantly contained TdT+ neurons reflecting the W-specific activity of these two hypothalamic structures. Consistent with our data, calcium imaging in vGluT2-cre mice showed that PVN glutamatergic neurons (including vasopressin, oxytocin, and CRF subtypes) are mainly active during W [63]. Opto-stimulation of these neurons or their projections to the lateral parabrachial nucleus/lateral septum rapidly awaken mice, while chemogenetic inhibition or lesions markedly reduced W amounts [12, 64, 65]. Activation of vasopressin PVN neurons also promoted W, an effect blocked by Orx antagonists when stimulating PVN terminals in the LHA [65, 66]. Together, these findings showed that the PVN is a critical structure for W expression, in particular through the activation of orexin-expressing neurons. The role of the SO nucleus in W has never been addressed and needs additional investigation.

In the LHA, approximately 20% of c-Fos+ PS-on neurons express MCH, Orx, or Lhx6. In the ZI, 11% of c-FoslJ neurons were positive for MCH or Lhx6. MCH neurons send extensive projections to different brain areas [67–71], including structures involved in regulating the sleep-wake cycle, stress- and anxiety-like behaviours, reward value of food consumption, body weight and energy balance [14, 19, 28, 31, 72–78]. In addition to the MCH peptide, some neurons co-express neurokinin 3 receptor (NK3R), the cocaine- and amphetamine-regulated transcript (CART) peptide [16, 79, 80], nesfatin-1 [81, 82] and neuropeptide-EI and GE [69, 83]. The existence of two subpopulations of MCH neurons with different embryological dates of origin, presence or absence of CART/NK3R expression, and different axonal projection patterns were demonstrated in both rats and mice [81, 84]. In rats, the first subpopulation which represents 2/3 of the whole population is located ventrally and medially and expresses CART/NK3R. These neurons predominantly project rostrally to the telencephalon. The second population lacks these markers and is located more dorsally and laterally with their axons projecting to the brainstem and spinal cord [85]. We show here for the first time that the ventromedial subpopulation of MCH neurons, are more activated during PSR, indicating that they could have different function. Additional studies are needed to confirm our results.

Previous studies in rats showed that only 2% of Orx neurons were c-Fos+ during PSR although the 150min PSR period included 23% of W [14]. In contrast, our results in mice show that 25% of Orx neurons were c-Fos+ during 120 min of PSR which included 19.4% of W while 44% were labelled with TdT during 2h of permanent W. Lee et al. [36] also using TRAP2-red mice reported that 32.5% of Orx+ neurons were activated during permanent W. Our results are not supported by juxtacellular unit recordings during W in rats and mice showing that Orx neurons virtually cease firing during SWS and display during PS only few phasic bursts synchronized to phasic muscular twitches (1-2 spikes). However, Orx neurons increased their firing a few seconds before the end of PS bouts followed by short arousal [6, 8]. However, it has been shown using calcium imaging that 7% of Orx neurons, which project to the pontine sublaterodorsal nucleus, are also active during PS [86]. In addition, using calcium imaging, Ito et al., [87] identified a subpopulation of Orx neurons (∼32%) that exhibit low but continuous activity during PS, unlike the majority of Orx neurons which are quiescent. From these data, it can by hypothesized that the numerous short W episodes occurring during PSR (22±1.9 episodes with a median of 16.5±5.4 seconds) could be responsible for our c-Fos staining. Further investigation, notably using calcium imaging is necessary to confirm this hypothesis.

We next assessed whether the activity of hypothalamic neurons expressing Meis2 might be linked to the sleep-wake cycle. To our knowledge, the mapping of neurons immunopositive for Meis2 has not been performed before. Our results are consistent with previous in-situ hybridization studies [47, 88, 89]. Meis2 contributes during development to the hypothalamic regionalization (tuberal, lateral, premamillary zones). There, it promotes GABAergic fate, specifies dopaminergic neurons during neuronal development (A13 in ZI) and acts as a key transcriptional factor [89–92]. It has been shown to be a specific marker of two GABAergic and one glutamatergic subpopulations of LHA neurons (among 15 of each type) separate from MCH, Orx and Lhx6 neuronal populations[24, 50, 93]. In Zi, Meis2 neurons were reported to be exclusively GABAergic [47, 89]. Here, we show for the first time that the rostral ZI, LHA, and AHA contain numerous Meis2+ neurons activated during PSR. Many of these neurons co-expressed c-Fos+ and, to a lesser extent, TdT+, whereas other hypothalamic nuclei depicted very few double-labelled cells. About one-fourth of c-Fos+ neurons in LHA and ZI were triple-labelled, compared to one-third of TdT+ neurons in LHA and half in ZI. These findings suggest that most Meis2 neurons in LHA and ZI are activated during PSR but not during W, contrasting with half of those activated during W which were also activated during PSR. In summary, our results show that Meis2 is expressed in previously uncharacterized hypothalamic subpopulations beyond Orx, MCH, and Lhx6 neurons. Further studies (e.g., calcium imaging, optogenetics, rabies tracing) are needed to define their role in sleep-wake regulation.

In conclusion, we show here that around 40% of neurons activated during PSR in LHA and ZI express at least one of the studied molecular markers (MCH, Lhx6, Orx or Meis2). Additional markers are needed for the remaining 60% of c-Fos+ neurons. Since it has been shown that most hypothalamic c-Fos+ neurons activated during PSR are GAD67-positive [15, 94, 95], these neurons should belong to one of the 13 subpopulations of GABAergic neurons identified in the LHA [24] not explored in the present report. Other hypothalamic regions, such as AHA and Tub also warrant further investigations, as they contain a high number of c-FoslJ neurons during PSR and may contain yet-unidentified neuronal populations involved in PS expression. Finally, our results importantly show that Meis2-expressing neurons located in ZI and AHA are specifically activated during PSR, implicating a potential a role in PS. It would be of great interest to study their activity using calcium imaging, opto- or chemogenetic approaches.

## Acknowledgements

This work was supported by CNRS (UMR5292), INSERM (U1028), University Lyon I and the French National Research Agency ANR-21-CE16-0030-01). Amarine Chancel received a one-year PhD grant from the Société Française de Recherche et Médecine du Sommeil (SFRMS). We thank the CIQLE Centre for Quantitative Imaging (SRF Lyon-Est Santé) for their help in scanning brain sections.

## Supplementary

We also calculated cell densities per mm2 by dividing the number of labelled neurons by the surface of the area considered, that is automatically calculated by the software (Figure Supp1).

**Cell density per mm^2^ = number of labelled neurons / area surface in mm^2^**

As illustrated in Supplementary Fig 1, small hypothalamic structures such as PVN and SO depicted the highest densities of both c-Fos+ or TdT+ neurons compared to the larger ones as LH, AHA and ZI endowed with the lowest densities of both cell types. In agreement with data expressed in raw numbers significantly higher densities of c-Fos+ than TdT+ neurons were also evidenced for AHA, Tub or ZI (*p<0.05) while PVN, RCh (*p<0.05) and SO (**p<0.01) showed higher densities of TdT+ than c-Fos+ neurons. Remaining hypothalamic structures such as Arc, DMH, PSTN, PHN, and VMH did not show statistical differences in cell densities of c-Fos+ and TdT+ neurons.

## Data availability

The data underlying this article will be shared on reasonable request to the corresponding author.

## Abbreviations

4-OHT: 4-hydroxytamoxifen;
AHA: anterior hypothalamus nucleus;
Arc: arcuate nucleus;
DMH: dorsomedial hypothalamus nucleus;
EEG: electroencephalogram;
EMG: electromyogram;
GABA: gamma-aminobutyric acid;
LHA: lateral hypothalamus area;
Lhx6: Lim homeobox 6;
MCH: melanin-concentrating hormone;
Meis2: Myeloid Ecotropic viral Integration Site 2 homeobox;
Orx: orexin/hypocretin;
PBS: Phosphate Buffered Saline;
PBST: Phosphate Buffered Saline with Triton- X100;
PFA: paraformaldehyde;
PHN: posterior hypothalamic nucleus;
PS: paradoxical sleep;
PSD: paradoxical sleep deprivation;
PSR: paradoxical sleep rebound;
PSTN: parasubthalamic nucleus;
PVN: paraventricular hypothalamic nucleus;
RCh: retrochiasmatic area;
SWS: slow-wave sleep;
SO: supraoptic nucleus;
TdT: TdTomato;
TRAP2: targeted recombination in active populations, 2^nd^ generation
Tub: tuberal nucleus;
VMH: ventromedial hypothalamus;
W: wakefulness;
ZI: zona incerta.

**Figure Supplementary 1. Comparison of the density of c-Fos+, TdT+ and double-labelled neurons in the 12 hypothalamic structures of interest**

Comparison of the density (mm²) of c-Fos+ and TdT+ and double-labelled neurons in 12 hypothalamic structures. AHA, ZI, Tub (*p<0.05) contain significatively more c-Fos+ than TdT+ neurons. PVN, RCh (*p<0.05) and SO (**p<0.01) contain significatively more TdT+ than c-Fos+ neurons (n=4 mice).

