## Supplementary figures and images for "Comparative distribution of the hypothalamic neurons activated during Wakefulness and Paradoxical (REM) sleep using TRAP2-red mice: contribution of Orexin, MCH, Lhx6 and a new marker Meis2"

### Supplementary figure 1

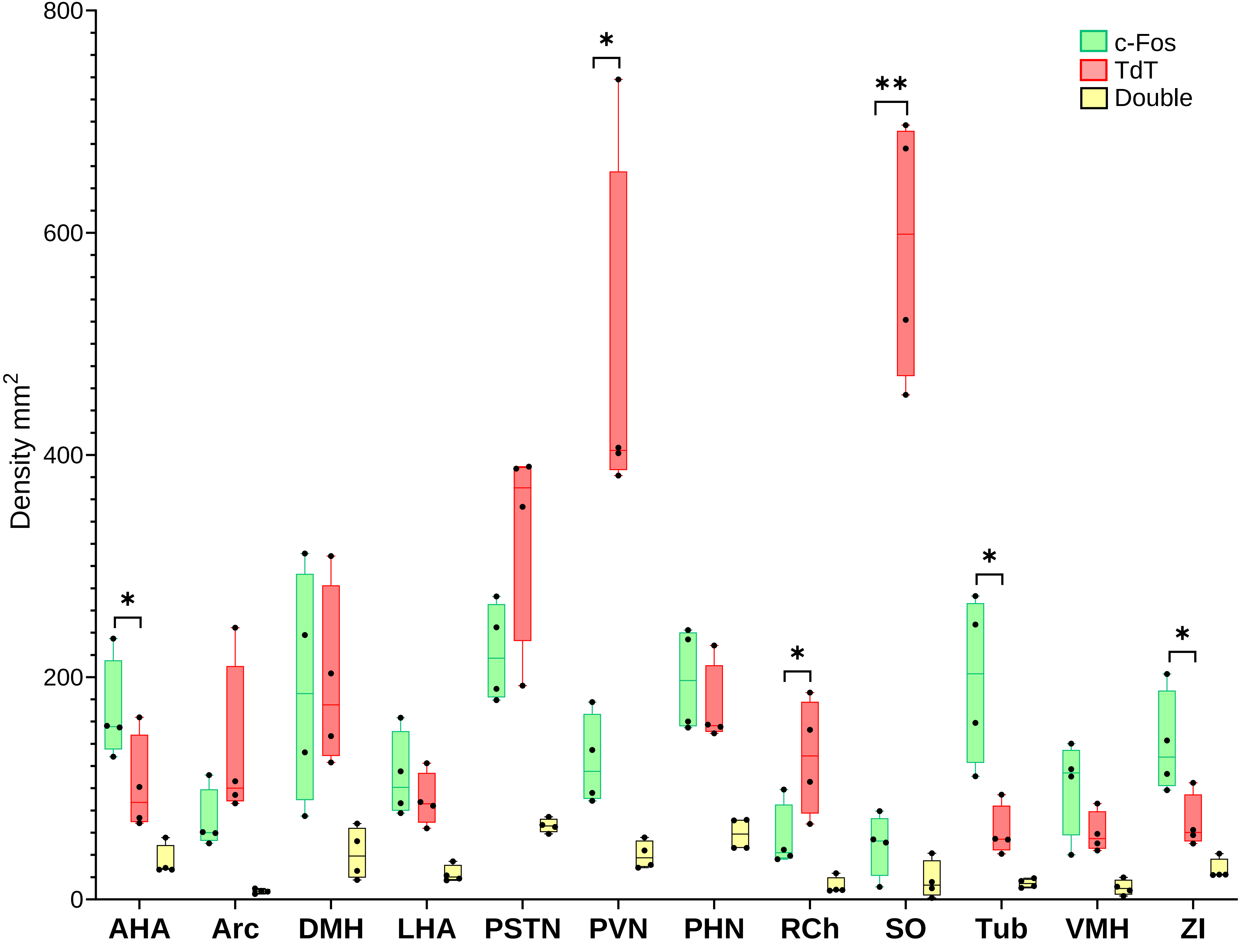
